# Structure of general-population antibody titer distributions to influenza A virus

**DOI:** 10.1101/090076

**Authors:** Nguyen Thi Duy Nhat, Stacy Todd, Erwin de Bruin, Tran Thi Nhu Thao, Nguyen Ha Thao Vy, Tran Minh Quan, Dao Nguyen Vinh, Janko van Beek, Pham Hong Anh, Ha Minh Lam, Nguyen Thanh Hung, Nguyen Thi Le Thanh, Huynh Le Anh Huy, Vo Thi Hong Ha, Stephen Baker, Guy E Thwaites, Nguyen Thi Nam Lien, Tran Thi Kim Hong, Jeremy Farrar, Cameron P Simmons, Nguyen Van Vinh Chau, Marion Koopmans, Maciej F Boni

## Abstract

Seroepidemiological studies aim to understand population-level exposure and immunity to infectious diseases. Results from serological assays are normally presented as binary outcomes describing the presence or absence of pathogen-specific antibody, despite the fact that many assays measure continuous quantities. A population’s natural distribution of antibody titers to an endemic infectious disease may in fact include information on multiple serological states – e.g. naiveté, recent infection, non-recent infection – depending on the disease in question and the acquisition and waning patterns of host immunity. In this study, we investigate a collection of 20,152 general-population serum samples from southern Vietnam collected between 2009 and 2013 from which we report antibody titers to the influenza virus HA1 protein using a continuous titer measurement from a protein microarray assay. We describe the distributions of antibody titers to subtypes 2009 H1N1 and H3N2. Using a model selection approach to fit mixture distributions, we show that 2009 H1N1 antibody titers fall into four titer subgroups and that H3N2 titers fall into three subgroups. For H1N1, our interpretation is that the two highest-titer subgroups correspond to recent infection and historical infection, which is consistent with 2009 pandemic attack rates. For H3N2, observations censored at the highest titer dilutions make similar interpretations difficult to validate.

## 1 Introduction

Influenza viruses circulate globally [1,2], with predictable wintertime epidemics in temperate zones and less predictable patterns of persistence and irregular epidemics in tropical areas [3]. The size or attack rate of an epidemic in any given area can be measured with basic seroepidemiological methods, by taking a cross-sectional sample after an epidemic period and testing for the presence or absence of virus-specific antibody. This approach is easily interpretable in a post-epidemic or post-pandemic period [4,5] as the seropositive individuals are those that were infected recently. However, in tropical countries, the timing of epidemics is irregular and difficult to predict [6], and analysis of cross-sectional serological data would need to take into account that samples may not have been collected during a period that follows an epidemic.

In influenza serology, the concentration of antibody in the blood directed against the hemagglutinin (HA) surface glycoprotein of the influenza virus is known to correlate positively with the immune status of the body [7–9]. The gold standard serological tests to detect the presence of antibody against a particular influenza virus are Haemagglutination Inhibition (HI) and Microneutralization (MN) [8,10,11], which give discrete readouts – from 10 to 2560 depending on the testing purposes. HI titers of 40 are generally considered protective [12,13], and HI titers of 20 or 40 are viewed to be indicative of past infection, but concerns about errors [14] and repeatability [13] have limited the ability of the HI assay to give accurate estimates of past infection.

A binary approach to serology has several drawbacks. The cutoff value for seropositivity is typically calibrated from a patient set of confirmed acute cases, by collecting convalescent serum samples a few weeks or a few months after infection. This means that the more appropriate application of the cut-off value is the identification of recent symptomatic infections rather than any past infections. Thus, applying the threshold approach to a population-wide dataset could lead to underestimation of the seroprevalence. Two other drawbacks of binary classification are that it reduces the information available to accurately describe the epidemiology of an endemic disease, and that it results in incorrect or inconclusive classifications for samples with borderline measurements [14-16].

Modern serological techniques developed over the past decade have aimed to address some of the shortcomings of HI and MN assays: *(i)* the large amount of serum required to test for presence of antibodies to multiple viruses, *(ii)* discrete titer readouts, and *(iii)* limited titer reproducibility and lack of standardization across laboratories. Improvements in these areas can be seen in a protein microarray (PA) assay developed by Koopmans et al [17]. The assay allows simultaneous serological testing for different influenza strains with as little as ten microliters of serum. This has helped increase the underrepresented contribution of infants and young children groups from whom large volumes of blood are not normally taken [18–24]. Additional advantages of the microarray include continuous and reproducible titer values and the ability to standardize and correct the results for inter-technician and inter-laboratory variations [17].

In general, the advantage of analyzing continuous serological measurements is that this results in better classification for borderline samples and more accurate estimation of the seroprevalence [15,25–28]. Statistical approaches using mixture distributions have proven successful in describing the uncertainty of individuals’ past infection status [26,27,29–34]. In some instances, mixture distribution approaches have shown that the population is best classified into three or more subgroups, and these subgroups are normally interpreted as having a different status of protection, past infection, and/or vaccination [15,27,32,34–41]. Note that in some of these past studies there was statistical evidence – using either Bayesian Information Criterion (BIC) or Akaike Information Criterion (AIC) – that the multimodal distribution fit better than unimodal and bimodal ones, but none of these studies took into account the epidemiological interpretations of the underlying process or performed visual inspection when choosing the best-fit model (see Rota et al [37] or Gay [42] for a typical analysis).

Recently, mixture distribution approaches have been extended to influenza serology by Steens et al [43] and te Beest et al [38,39], in which post-pandemic data were collected right after the first wave of the H1N1 Pandemic 2009 in the Netherlands and analyzed by protein microarray. In these studies, mixture analysis was meant to measure the proportion of recent infection of a novel virus in the population. In the analysis presented here, we consider cross-sectional seroepidemiological data collected from southern Vietnam, using continuous titer measurements via the PA method. As there is very little influenza vaccination in Vietnam (lower than 1% annual coverage according to commercial vaccine sales data), these results are intended to provide general insights into long-term patterns of influenza circulation.

## 2 Methods

Residual serum samples were collected from four hospital laboratories in southern Vietnam: the Hospital for Tropical Diseases in Ho Chi Minh City (urban, densely population), Khanh Hoa Provincial Hospital in Nha Trang city (small urban, central coast), Dak Lak Provincial Hospital in Buon Ma Thuot city (central highlands, rural), and Hue Central hospital in Hue City (small urban, central coast). Samples were collected from July 2009 to December 2013 on a bimonthly basis; 200 were included in each collection from all age groups (neonates to elderly individuals in their 90s). Samples were anonymized, delinked, and labeled with age, gender, originating hospital ward (HIV wards were excluded), and date of collection. Samples were collected from both inpatients and outpatients and are believed to represent the hospital-going population in their respective cities. The sample collection described here is part of a large ongoing study in serial seroepidemiology [44] aimed at describing the dynamics of influenza circulation in southern Vietnam. The study was approved by the Scientific and Ethical Committee of the Hospital for Tropical Diseases in Ho Chi Minh city and the Oxford Tropical Research Ethics Committee at the University of Oxford.

The samples were tested for presence of influenza antibodies using a protein-microarray (PA) method [17], at serial four-fold dilutions from 20 to 1280, to test for IgG antibody to the HA1 component of 16 different influenza viruses [44]. Two-fold dilutions were used in some instances; see validation of this approach in Appendix Section 2. A sample of the international standard (IS) for testing antibody response to influenza A H1N1 Pandemic 2009 (H1-09) was included on every slide to correct for inter-laboratory, inter-technician, and inter-slides variations [17] (Appendix Section 1.2). Assay repeatability was assessed using a positive control and replicates of patient samples (Appendix Section 3). Titers were defined as the dilution at which samples yield a median response between the minimum and maximum luminescence values of 3000 and 65355. Titers of all human samples on each slide are normalized based on the IS titers of the reference antigen against its geometric mean (Table S2). In this analysis, titers to the 2009 H1N1 virus (A/California/6/2009) and recently circulating H3N2 viruses (geometric mean titer to A/Victoria/210/2009 and A/Victoria/361/2011) were analyzed.

To describe the distribution of influenza antibody titers in the Vietnamese population, titer values were separated by site, adjusted to their province’s age and gender distribution [45] (Appendix Section 4), and plotted as a simple weighted histogram (Figure 1). A series of mixture models was used to fit this distribution, with the assumption being that individual samples have one of several immune statuses which are represented by the different components in the mixture model. Our hypothesis was that the sample population consists of different subpopulations with different antibody levels depending on their infection history and that each of these components could be represented by a single parametric distribution.

**Figure 1:**
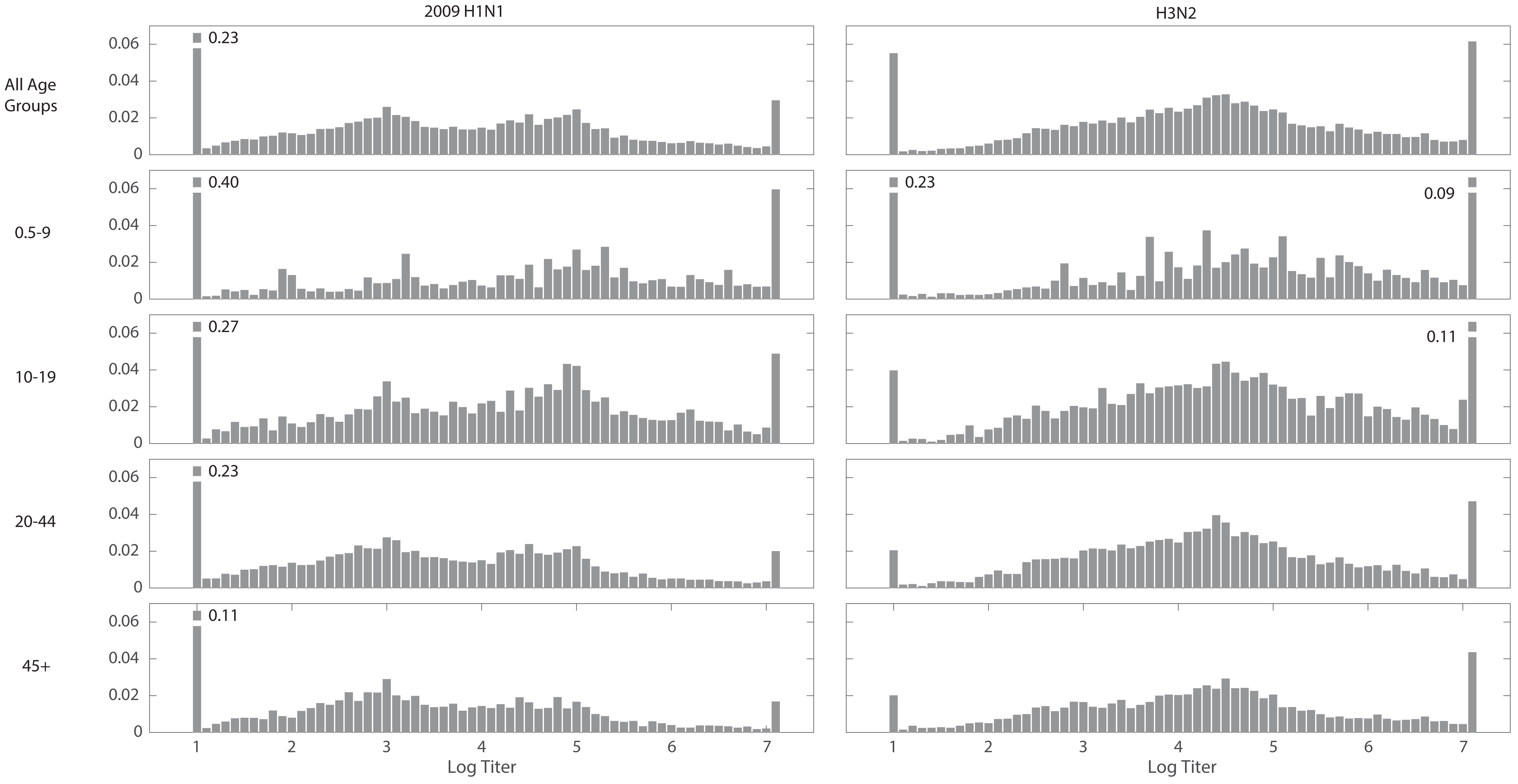
Antibody titer histograms for n = 20,152 individuals, plotted for all ages (top panels) and by age group (bottom four panels). Titers shown are to the HA1 components of the 2009 H1N1 pandemic influenza virus (left column) and to recently circulating H3N2 viruses (right column). The fractions of individuals with titers below the detection limit of 20 and above 1280 that were out of the plotting ranges are given next to the respective bar. Histograms were weighted to adjust for age and gender according to the Vietnam national housing census in 2009 for the four collection sites.

Titers were log-transformed and assumed to come from a *C*-component mixture distribution with the corresponding likelihood:

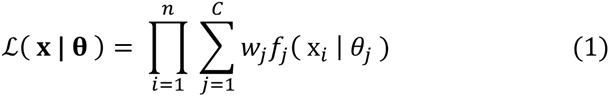

where *f* is the probability density function of a normal distribution with parameters *θ*_*j*_ and **w** = (*w*_1_, *w*_2_, …, *w*_*C*_) is the vector of component weights in the mixture. The log-likelihood was defined as:

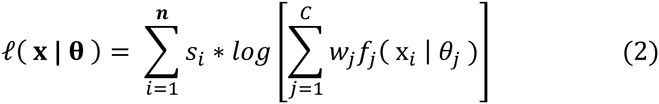

in which the *s*_*i*_ parameters are sampling corrections to adjust the sample age and sex distribution to the population’s true demographic distribution; *f*_*j*_(*x*_*t*_ | *θ*_*j*_), j = 1, 2,.., *C* is the probability density function that a given sample *x*_*i*_ belongs to the *j*th-component in the mixture. *C* is the number of mixture components [46,47].

The microarray assay produces continuous log-titer results between 1.0 (titer of 20) and 7.0 (titer of 1280). To account for these detection limits, an extra probability weight *w*_0_ was added at 20 to account for samples that had antibody concentrations at or below the detection limit of 20. This can be considered a zero-inflated mixture model, where titers of 20 are the “zeroes”. With this added probability mass, the continuous component distribution must be discretized into probability mass functions; hence the distributions *f* formally represents discretized versions of continuous density functions (Appendix Section 5). At the upper detection limit of 7.0, the mixture distribution was censored assuming that individuals with titers of 7.0 represented a class of seropositive individuals with a real titer value if the assays had been continued to be diluted until the real titer was found. Censoring on the right and truncating on the left gave the best fit among the four combinations.

Thus, the likelihood in (1) was modified as:

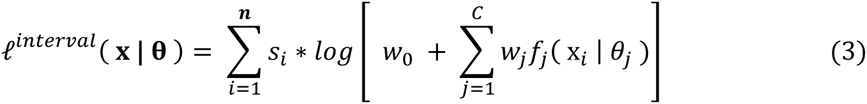

Maximum likelihood estimation was carried out using the Nelder-Meade algorithm implemented in Java 8.0 (Apache Commons Math 3.3). Global optima and convergence were assessed by starting searches from different sets of the initial conditions. Weibull, Gamma, and normal distributional forms were tested for the mixture components, and as there was little difference in the fits (with Weibull distributions having slightly worse fits; not shown), normal distributions were chosen for the analysis.

For multi-component mixture models, the likelihood ratio test between a specific model and its immediate predecessor (e.g. *n* components versus *n*-1 components) is not a valid statistical comparison. Since interchanging the components’ identity gives the same mixture likelihood [47], the regularity conditions do not hold for the likelihood ratio test to have its usual *χ*^2^ distribution. Thus, the most appropriate number of mixture components was chosen (1) by Bayesian Information Criterion to take into account the number of samples, and (2) by a qualitative inspection of the means and variances of the components to ensure that (*a*) multiple means did not overlap and (*b*) variances and weights were not too small, which would make them not epidemiologically meaningful.

## 3 Results

A total of 20,152 sera were collected and tested for antibody concentrations by protein microarray. The samples represent patients attending hospitals in four cities – Ho Chi Minh City (n=5788), Nha Trang (n=5630), Buon Ma Thuot (n=4144), and Hue (n=4590). Titer distributions varied by age, as expected (Figure 1) but did not vary by site (Figure S6 and S7). Figure 1 shows the age-stratified titer distributions to the HA1 component of the 2009 H1N1 virus and the most recently circulating H3N2 variants. If individuals fall neatly into seropositive (exposed) and seronegative (unexposed or naïve) categories, a mixture model of two components would classify samples into two subgroups, which clearly was not the case as a broad range of titers was observed for both subtypes across all age groups. Thus, a mixture distribution fitting approach was employed to determine the appropriate number of mixture components necessary to accurately describe the titer data.

To identify the appropriate number of subgroups in the titer distributions, the Bayesian Information Criterion (BIC) was used to select the number of components in the mixture. Figure 2 shows six rows of distribution fits – from single-component fits to six-component fits – for antibody titers to the 2009 H1N1 pandemic virus; data are shown both aggregated and by site (Figure S8 for H3N2). The BIC is shown in the upper-right corner of each panel, and the BIC improvement from *n* components to *n + 1* components varied depending on the number of samples in the dataset; see Table 1 for 2009 H1N1 and Table S4 for H3N2. For both subtypes, it is clear that a binary classification of titer is not the best interpretation of the natural antibody titer distribution, as the one- and two-component models (top two rows) did not capture the underlying structure of the dataset adequately. When stratifying the data by site (sample size ∼4,000), the BIC consistently selected four components as the best model for the H1N1 data (five for Hue, but weakly: ΔBIC=18) and three components as the best model for H3N2.

**Table 1.** Change in BIC scores as the number of normal distributions in the mixture increases from one to six for 2009 H1N1, for the aggregated data as well as the individual collection sites. The values of the negative sum of the log likelihood were weighted to adjust for age and gender according to the Vietnam national housing census in 2009. The first row shows the exact values of sum of negative log-likelihood (SumNegLLH). Bold numbers represent the mixtures with the best BIC.

|  |  | ALL SITES<br>(N = 19335) | HUE<br>(N = 4390) | KHANH HOA<br>(N = 5053) | HCMC<br>(N = 5753) | DAK LAK<br>(N = 4139) |
| --- | --- | --- | --- | --- | --- | --- |
| SumNegLLH<br>(C=1) |  | 71504.04 | 15812.88 | 18726.69 | 21496.01 | 15432.06 |
| ΔBIC | 1 to 2 | -826.40 | -170.79 | -182.44 | -215.33 | -208.86 |
|  | 2 to 3 | -410.07 | -74.55 | -79.83 | -137.95 | -131.34 |
|  | 3 to 4 | -318.12 | -66.76 | **-56.37** | **-77.14** | **-32.68** |
|  | 4 to 5 | -19.87 | **-18.32** | 8.68 | 16.82 | 6.10 |
|  | 5 to 6 | **-27.42** | 2.13 | 24.23 | 9.57 | -2.59 |

**Figure 2:**
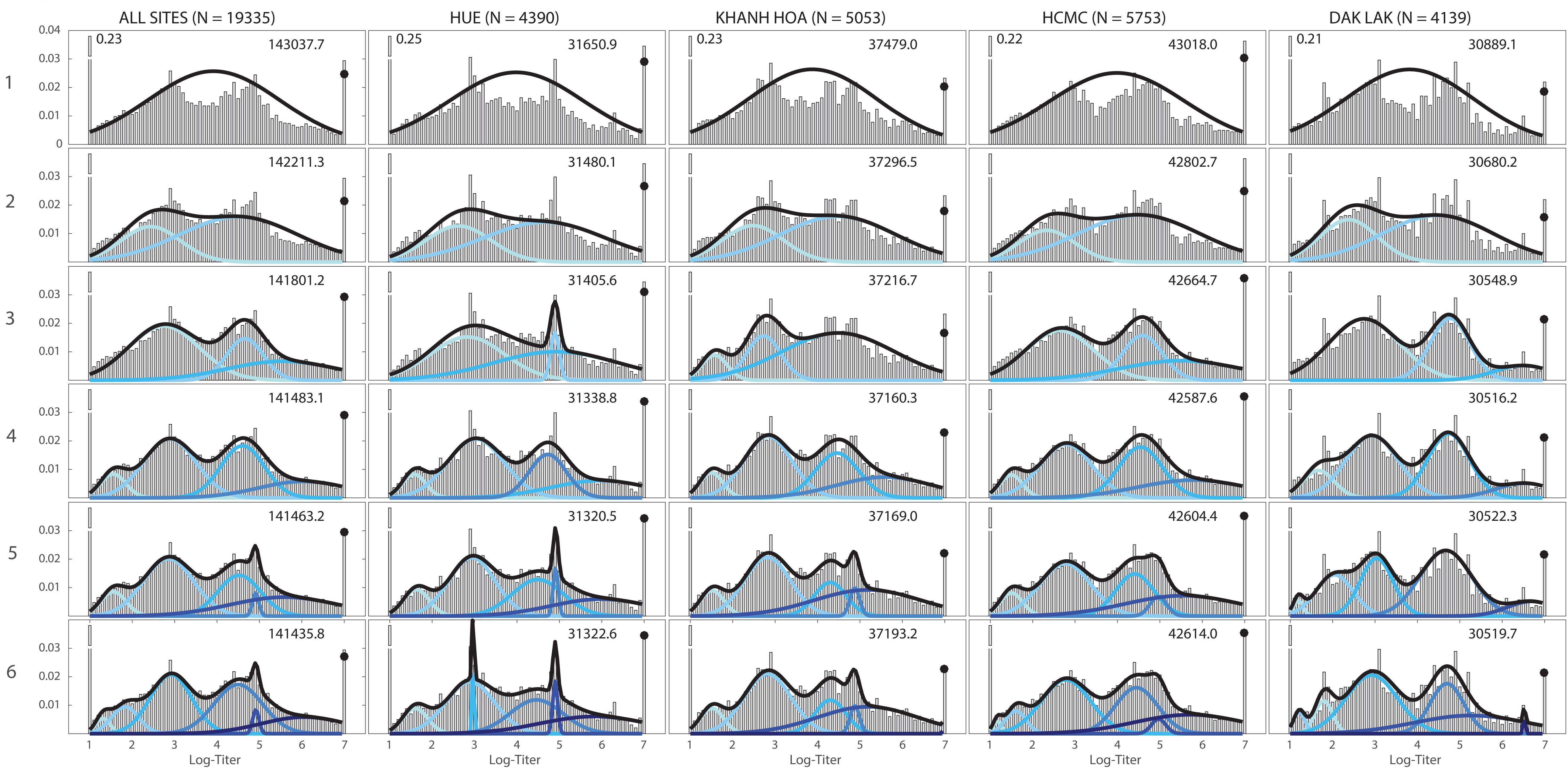
Titer histograms for 2009 H1N1, showing fit results for mixture models with different numbers of normal components (top to bottom; the label to the left of the *y*-axis is the number of mixture components) and grouped by collection sites. Histograms are weighted to adjust for age and gender according to the Vietnam national housing census in 2009 for each of the four collection sites. The blue lines in each panel are the normalized probability density functions of the component distributions with darker colors used for increasing *μ*. The black lines show the full mixture distribution density, and the black dots are the estimated cumulative distribution of the mixture models at 7.0 (titer of 1280). The numbers in the upper right corner of each panel are the BIC scores of the model fits. The fractions of individuals with titers below the detection limit of 20 and above 1280 that were out of the plotting ranges were given next to their respective bars.

Interpreting the mixture distributions in the context of the known seroepidemiology of influenza suggests that the three component models and four component models are the best descriptions of the populations’ antibody titer distribution. The five- and six-component models either overfit the data (according to the BIC) or included low-variance/low-weight components, which would correspond to an implausible population subgroup with a very specific antibody titer (Figure 3). This was readily seen in the aggregate data which is why the BIC-selected models of the by-site data (∼4000 data points each) are likely to be better explanations of the structure of these titer distributions. For H1N1, the three- and four-component models identified similar titer subgroups with the four component models giving better BIC. Going from three to four components did not significantly affect the last two components (weights, means, and standard deviations). The main difference between these models was the presence of the first small peak (titer range 19.7 to 45.0) which helped improve the BIC and minimized the overlap between the two highest titer components. Epidemiologically, this first peak represents seronegative individuals with very low titer values [38,39,48]. For H3N2, the results were clearer as the BIC selected three components as the best model for all sites (Figure S8).

**Figure 3:**
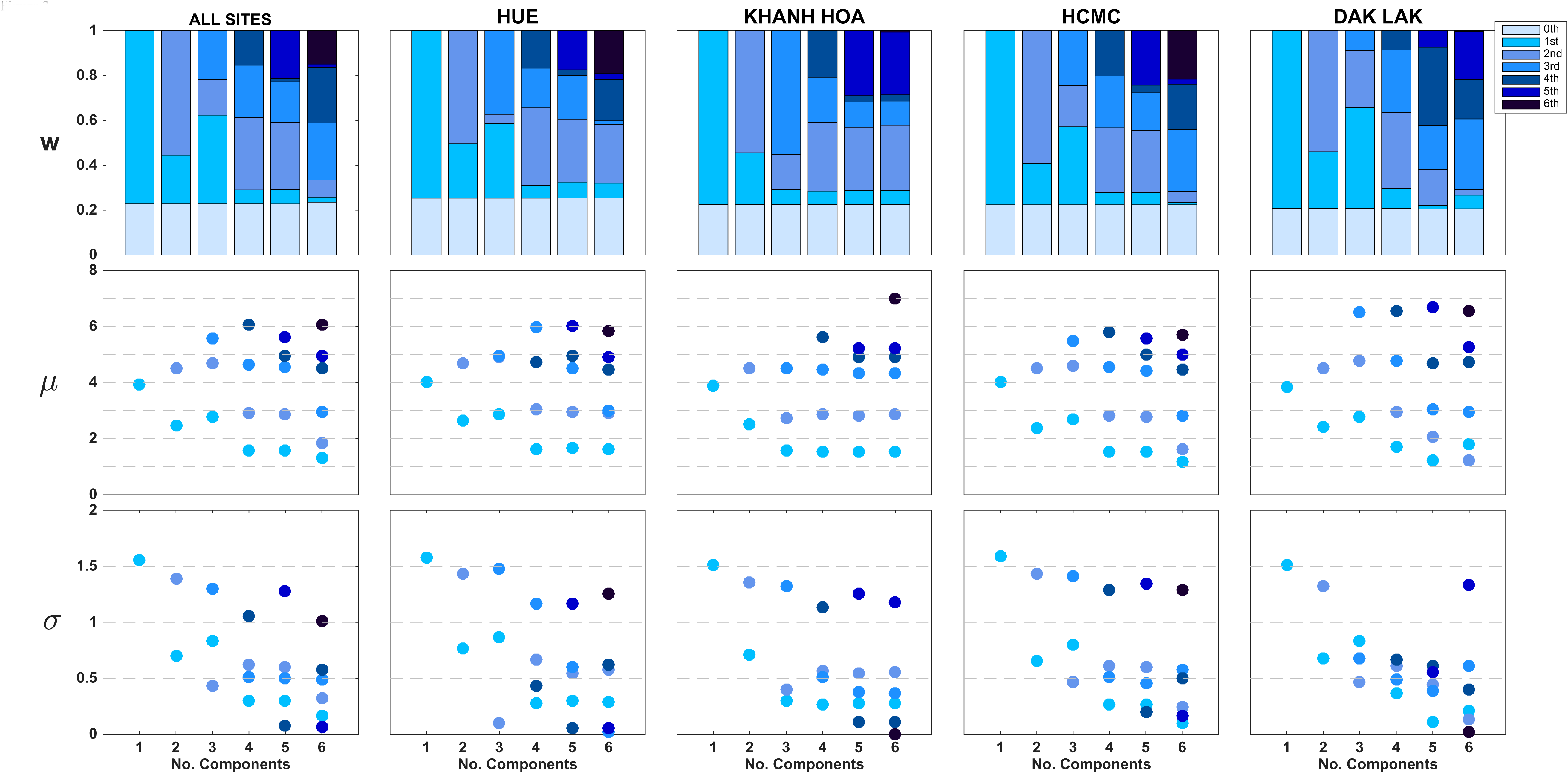
Visualization of model selection process for 2009 H1N1 titer-distribution models from Figure 2. The *y*-axes show the fitted values of *w*_*i*_ (mixture weights), *μ*_i_ (means), and *σ*_i_ (standard deviations). Components’ shades were ranked from lightest to darkest in the order of increasing *μ*. In the top panel, the “0^th^ component” represents the point mass w_0_ placed at 20 for titers below the lower detection limit of 20. Note that in many cases for five or six components, the weights or standard deviation parameters are close to zero; for some cases, two of the inferred mean parameters are very close to each other.

The three- and four-component mixtures indicate that these data can be used to develop a more informative serological classification for influenza. Using known results for this microarray assay [38,43], titers below 100 would be classified as negative or ‘not previously exposed to this particular influenza strain’. For H1N1, this means that that the first component (μ_1_ = 30) and the second component (μ_2_ = 75) would correspond to negative individuals. Similarly for H3N2, negative individuals would be represented by the first component (μ_1_ = 80). The second-highest titer component had mean μ_3_=247.2 (95% CI 240.0−253.7) for H1N1 and μ_2_=213.3 (95% CI 204.8−221.5) for H3N2. The highest titer component has mean μ_4_=671.1 (95% CI 484.7–818.6) for H1N1 and μ_3_=455.0 (95% CI 401.4−536.7) for H3N2. Confidence intervals were computed using likelihood profiles [47]. The natural interpretation of these high titer subgroups is that they represent different times since last infection. As it is known that the influenza antibody decay rate is fast enough to be observed in the first six to twelve months after an acute infection [49,50], for H1N1 the highest titer subgroup may be an approximate designation for recently infected individuals, and the second highest titer subgroup may correspond to ‘historically infected’ individuals, i.e. individuals infected at some point in the non-recent past.

For H1N1, these interpretations were able to be validated using post-pandemic sera, and an ROC curve was constructed using the methods presented in te Beest et al [38] (Figure 4). Assuming that the highest-titer component (*w*_4_) of the mixture distribution corresponds to recently infected individuals and the second highest-titer component (*w*_3_) corresponds to historic infection, one would expect to be able to use the weights *w*_3_ and *w*_4_ as proxies for the pandemic attack rate. Looking at samples collected from January 2010 to June 2010 – i.e. after the first wave of the 2009 influenza pandemic in Vietnam [23,51] – the proportions of individuals that were recently infected with 2009 H1N1 were highest among younger individuals (0.14, 0.23, 0.08, and 0.16, for the 0.5–9,10–19, 20–44, and ≥45 age groups, respectively), while the proportions of historically-infected individuals were approximately equal among age groups (0.16, 0.22, 0.23, and 0.20, for the same age groups). The estimates of 14% of children aged 0.5–9 and 23% of children aged 10-19 falling into the recently infected category will be slight underestimates of pandemic attack rate as the post-pandemic sample here includes samples collected through June 2010. Nevertheless, these are within the expected ranges of the attack rate of the first year of the 2009 pandemic. For older individuals, pandemic attack rates are more difficult to validate but it is important to remember that older individuals had measurable antibody titers to 2009 H1N1 prior to the arrival of the new pandemic virus [16].

**Figure 4:**
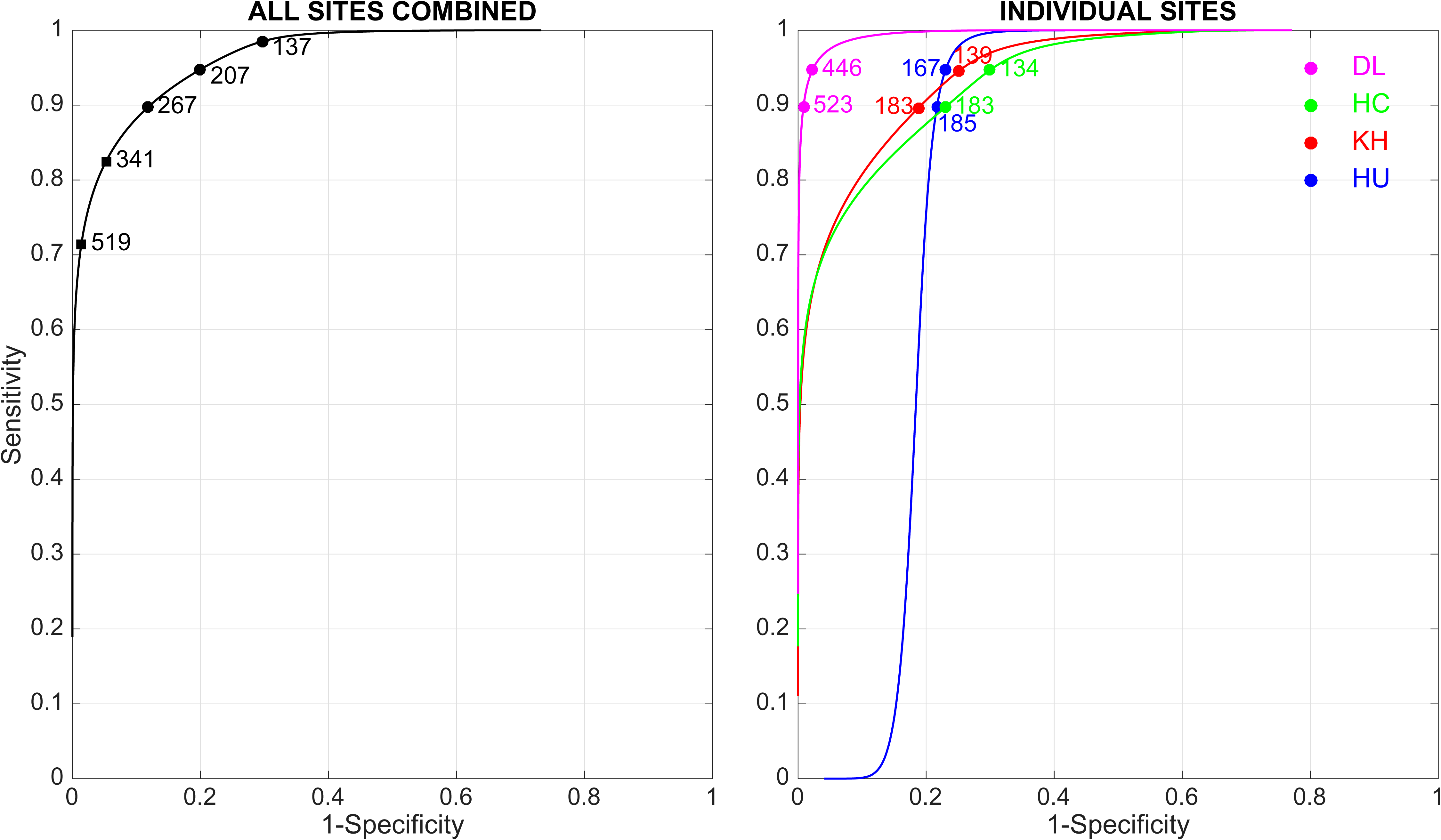
ROC curves for H1N1 using four mixture components, presented as in te Beest et al [38]. A titer cut-off is determined to have a particular sensitivity and specificity based on how well it sorts individuals into the different mixture components, with the fourth component representing positivity. In the left panel, the mixture distribution for all sites is used. The titers indicated on the curve show, from left to right, 99% specificity, 95% specificity, 90% sensitivity, 95% sensitivity, and 99% sensitivity. In the right panel, mixture distributions for individual sites are used. The indicated titers show 90% sensitivity and 95% sensitivity. Abbreviations: DL, Dak Lak; HC, Ho Chi Minh City; KH, Khanh Hoa; HU, Hue.

For H3N2, the best-fit mixture models had larger variances than the best-fit models for H1N1. The log-titer ranges (±2σ) for the three H3N2 titer groups were 26–240, 114–456, and 68–3045. Thus, the discriminatory power between the last two components was not as good as for H1N1 (Figure S8). One plausible explanation is the existence of an additional fourth peak for the H3N2 titers describing individuals with titers above the upper limit of detection (≥1280). In our sample set, the proportions of individuals with H3 titers equal to 1280 were two to three times higher than those for H1N1 in the same age category. The large standard deviations of the last component for H3N2 may have been the result of the high fraction of right-censored samples with titers ≥1280.

Thus, the epidemiological interpretation of the H3N2 mixture components cannot be validated at present. Using all samples, the proportions of individuals in the highest titer group (third component) are 0.46 (ages 0.5–9), 0.49 (ages 10–19), 0.43 (ages 20–44), and 0.34 (≥45). These are unlikely to represent recent attack rates of H3N2 epidemics and are more likely to represent historical infection, i.e. individuals who have been exposed to the currently circulating H3N2 strain. This is consistent with our conjecture that a fourth peak in the H3 titer distribution may not be visible due to truncation at a titer of 1280, but we were not able to confirm this with the current data as the samples were not diluted past 1:1280.

For both subtypes, the individual components in the mixture models did not correspond to any particular age groups, and stratifying the samples by age did not explain any particular component of the mixture (Figure 5 for H1N1 and Figure S10 for H3N2). All age groups included individuals with high, medium, and low titer levels.

**Figure 5:**
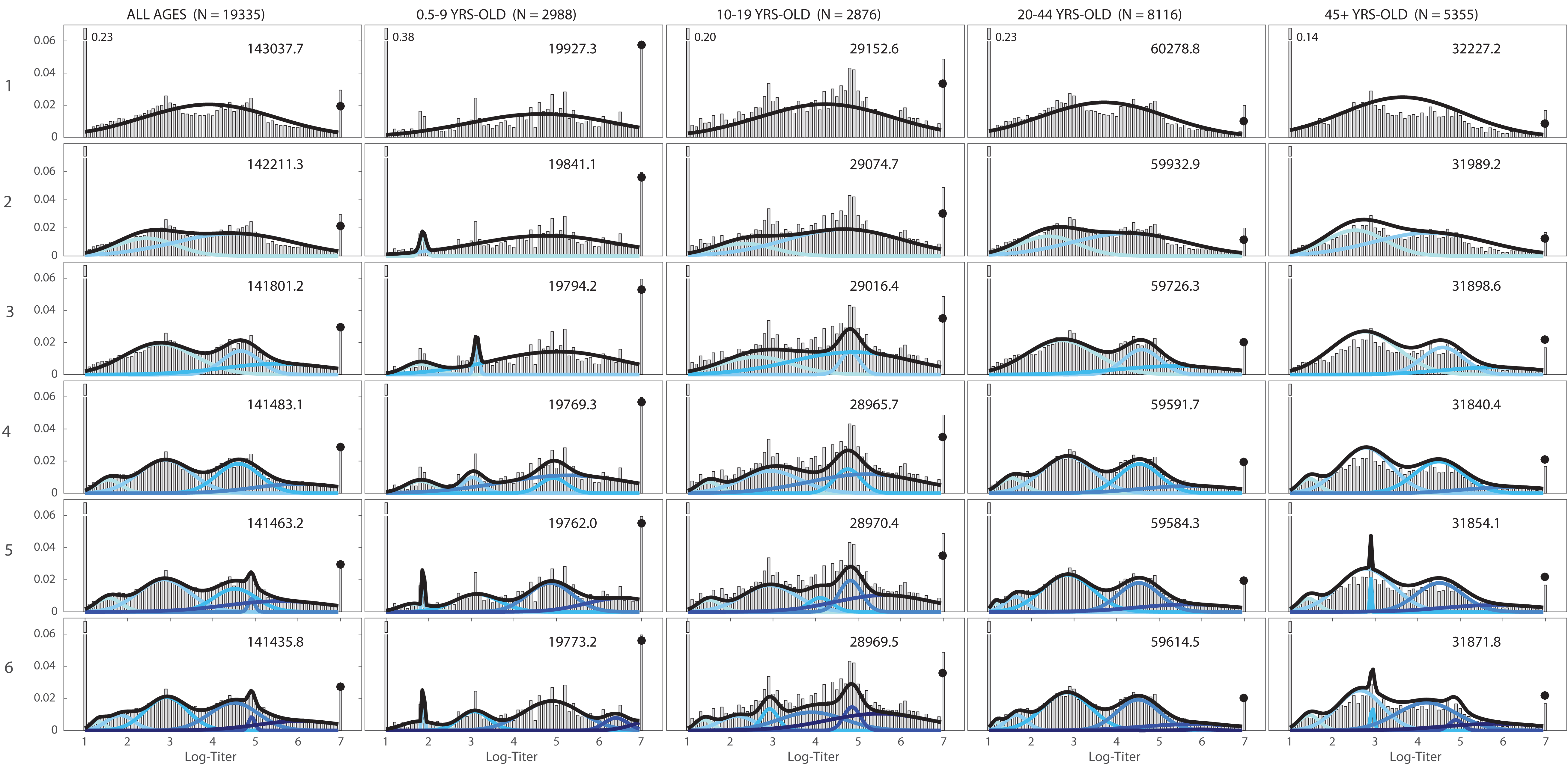
Titer histograms and fit results for mixture models with different numbers of components (label on the left is the number of mixture components) and grouped by different age groups recommended by the CONCISE (http://consise.tghn.org/) for 2009 H1N1 influenza. Histograms are weighted to adjust for age and gender according to the Vietnam national housing census in 2009. The numbers in the upper right corner of each panel are the fitted BIC scores of the respective model. For each panel, the blue lines are the normalized probability density of the component distributions with darker colors used for increasing *μ*. Black lines are the total mixture distribution density; and the black dots are estimated probability weight of the mixture model for titers ≥ 7.0. The fractions of individuals with titers below the detection limit of 20 and above 1280 that were out of the plotting ranges are shown right next to their respective bars.

## 4 Discussion

Using a large collection of serum samples and a continuous measurement of antibody titer, we were able to describe the natural distribution of antibody titers to the 2009 H1N1 and H3N2 subtypes of influenza virus. As there is almost no influenza vaccination in Vietnam and as influenza in Vietnam is characterized by a combination of local persistence and annual/binannual outbreaks [52–54], characterization of titer distribution in this context is a useful general approach for looking at the immune status of a population at equilibrium with an endemic infectious disease. With a mixture model approach, we were able to identify the presence of multiple exposure groups in the population according to their titers. Our interpretation of these multiple exposure groups – according to titers measured for confirmed cases [48] and past measurements of the rate of antibody waning [49,50] – is that they represent recently infected individuals, historically (i.e. not recently) infected individuals, and naïve individuals. Note that for influenza, a naïve individual is one who has not been exposed to the currently circulating strain, which means that there will be naïve individuals in all age groups.

A mixture distribution approach does not guarantee that individuals can be easily classified into one of several titer subgroups. With substantial overlap in the mixture components, individuals typically have partial membership in two, sometimes three, titer subgroups. In addition, individual variation will have a large effect on titer interpretations. A high-titer sample could represent a recent infection, but individuals can maintain high titers longer than the mean duration observed in clinical studies. This would normally, but not exclusively, be observed in children. Likewise, lower antibody titers (in the 200–250 range) could indicate historical past infection, a low response to a recent infection [55], or a recent but mild infection. With serological data alone, these scenarios cannot be distinguished. For subtype H3N2 specifically, low titer levels could indicate cross-reactions between antibodies generated to an older influenza variant and the recent H3N2 HA1 proteins spotted on the protein microarray.

An important new question generated by this study is why H1N1 titer distributions appear bimodal but H3N2 titer distributions appear unimodal. One possibility is that the censoring of H3 titers at 1280 truncated the true bimodal appearance of the H3 titer distribution (see Figure S11). A second possibility is that some feature of the infection, immunogenicity, or waning process for H1N1 operates differently on different segments of the population. An important difference between these two subtypes is that H3 infections lead to higher antibody titers (as measured in HI and MN assays) than H1 infections, and for this reason the truncation hypothesis may be the more plausible one. A second difference lies in the subtypes’ lineage history, which suggests that separating the samples into H1N1 lineage-exposure groups (pre-1957, post-1977, post-2009) may account for the bimodal pattern in H1N1. However, separating the samples by birth year (0.5-50 years-old, and ≥60 years-old) did not provide any evidence for this effect (Figure S12 and Table S6).

A major challenge in influenza seroepidemiology is that it is difficult to take into account the effects of original antigen sin [56,57] or age-dependent seroconversion (ADS). Age-dependent seroconversion is distinct from original antigenic sin in that ADS assumes that individuals of different ages seroconvert to different titer levels irrespective of the individual’s infection history. In principle, the effect of ADS should be detectable for 2009 H1N1 infections in individuals younger than 50, as for these individuals an exposure to the 2009 virus would have been a first exposure. However, the component distribution means (*μ*_*i*_ parameters) and the component weights (*w*_*i*_) are not separately identifiable in the mixture model. Thus, we cannot state that the ‘recently infected’ titer subgroups are comparable across age groups, as the inferential process will make the exact definition of recency different for the 10-19 age group than for the 20-44 age group. Even if we were to assume that the fourth mixture components should be comparable across age groups, the titer means denoted by *μ*_4_ in Figure 4 do differ but are within one standard deviation of one another. Thus, there is a lack of evidence for ADS in our titer data. As we only considered recent antigens in this analysis, effects of original antigenic sin were not considered.

With a clinical study near completion, we will soon be able to validate the titer interpretations obtained from the mixture-distribution approach presented here. As has been found in other recent studies [58,59], the waning rate of influenza-specific IgG antibodies is crucial to interpreting antibody titers measured in serological cross-sections. Depending on the titer cutoff chosen or the waning rate used, measured seropositivity in a population cross-section could represent any range of infection history, from recent infection to an older historical infection. The next critical step in this analysis will be using titer data from follow-up on confirmed cases to determine if the natural distribution of antibody titers should conform to the recent, historical, and naïve categories as presented here.

## Acknowledgements

This work was funded by the Wellcome Trust Grants 089276/B/09/7 (NTDT, DNV, PHA, HML, HLAH, GET, JF, NVVC), 097465/B/11/Z (ST), 098511/Z/12/Z (TTNT, NHTV, NTH, NTLT, MFB), and by a British Medical Association HC Roscoe Award (2011-2014). CPS is funded by the National Health and Medical Research Council of Australia. MK, EdB, and JvB are funded by Dutch Ministry of Economic Affairs, Agriculture, and Innovation, Castellum Project.

### Author Contributions

NTDN, JF, CPS, NVVC, MK, and MFB conceived the study. NTDN performed all the analysis and wrote the first draft of the paper. NTDN and MFB validated all analyses and co-wrote the paper. SB, GET, and CPS edited the paper. EdB printed and validated the microarrays. TTNT, NHTV, and PHA performed all of the laboratory assays. ST and DNV contributed conceptually to determining how titer dynamics should behave in an endemic setting. JvB wrote the initial script for positive control correction. TMQ and HML analyzed fluorescence signals. NTH, NTLT, HLAH, VTHH, NTNL, TTKH coordinated sample collection at four sites over five years. All authors read and approved the final results and interpretations in the article.

